# Noninvasive lung cancer detection via pulmonary protease profiling

**DOI:** 10.1101/495259

**Authors:** Jesse Kirkpatrick, Andrew D. Warren, Tuomas Tammela, Peter M. K. Westcott, Justin C. Voog, Tyler Jacks, Sangeeta N. Bhatia

**Author notes:** These authors contributed equally to this work (Jesse Kirkpatrick, Andrew D. Warren).

## Abstract

Lung cancer is the leading cause of cancer-related death and patients most commonly present with incurable metastatic disease. National guidelines recommend screening for high-risk patients with low-dose computed tomography (LDCT), but this approach has limitations including high false positive rates. Activity-based nanosensors (ABNs) detect dysregulated proteases *in vivo* and release a reporter to provide a urinary readout of disease activity. Here, we demonstrate the translational potential of ABNs by coupling ABN multiplexing with intrapulmonary delivery to detect early-stage lung cancer in an immunocompetent, genetically engineered mouse model (GEMM). The design of the multiplexed panel of sensors was informed by comparative transcriptomic analysis of human and mouse lung adenocarcinoma data sets and *in vitro* cleavage assays with recombinant candidate proteases. When employed in a *Kras* and *Trp53* mutant lung adenocarcinoma mouse model, this approach confirmed the role of metalloproteases in lung cancer and enabled accurate early detection of disease, with 92% sensitivity and 100% specificity.

## Introduction

Lung cancer is the most common cause of cancer-related death (25.3% of cancer deaths in the U.S), with dismal 18.6% five-year survival rates^1^. Key to this high mortality is the fact that 57% of lung cancer patients have distant spread of disease at the time of diagnosis^1^. Because patients with regional or localized disease have six-to 13-fold higher five-year survival rates than patients with distantly spread disease^1^, significant effort has been dedicated to improving diagnostic sensitivity. Screening with low-dose computed tomography (LDCT) is recommended in high-risk patients (adults aged 55 to 80 with a 30 pack-year smoking history^2^) and enables a relative reduction in mortality of 20% when compared to the previous standard, chest radiography^3^. However, these screening tests are expensive^4^, have high false positive rates (~96%^3^) and potentially expose patients to biopsy-related complications, raising concern for overdiagnosis and increased healthcare-associated cost burden^5,6^.

Great strides in the field of molecular diagnostics have yielded promising approaches that may be used in conjunction with LDCT for lung cancer screening. Circulating tumor DNA (ctDNA) has emerged as a promising tool for noninvasive molecular profiling of lung cancer^7–10^. However, the presence of ctDNA has been shown to scale with tumor burden and there are fundamental sensitivity limits for early stage disease^7,10,11^. To achieve high-sensitivity detection of ctDNA in stage I-II cancer patients, it is estimated that large (>80 mL) blood volumes would be needed with current methodologies, potentially limiting the widespread adoption of this approach^12^. Similarly, circulating tumor cells (CTCs) may be detected in patients with advanced-stage non-small cell lung cancer (NSCLC), but the sensitivity of CTCs for detection of nonmetastatic disease remains low at present^13–16^. Finally, transcriptional profiling of bronchial brushings can enhance the diagnostic sensitivity of bronchoscopy alone, even for peripheral and early-stage pulmonary lesions, an approach that leverages the “field of injury” that results from smoking and other environmental exposures^6,17^. However, as with any invasive procedure, bronchoscopy carries the risk of attendant complications such as pneumothorax^18^.

Rather than relying on imaging techniques or detection of endogenous biomarkers in circulation, we have developed a class of “activity-based nanosensors” (ABNs) that monitor for a disease state by detecting and amplifying activity of aberrant proteases to generate urinary reporters^19–24^. Protease activity is dysregulated in cancer, and proteases across catalytic classes play a direct role in all of the hallmarks of cancer, including tumor growth, angiogenesis, invasion, and metastasis^25–30^. ABNs leverage dysregulated protease activity to overcome the insensitivity of previous biomarker assays, amplifying disease-associated signals generated in the tumor microenvironment and providing a highly concentrated urine-based readout. We have previously explored the sensitivity of this approach via mathematical modeling^31^ and cell transplant models^23^. However, to drive accurate diagnosis in a heterogeneous disease, a diagnostic must also be highly specific. Here, we explore the potential to attain both sensitive and specific early disease detection through multiplexing of 14 ABNs in an immunocompetent GEMM, which better recapitulates key aspects of human disease and allows for evaluation of diagnostic accuracy at the earliest stages of tumorigenesis. To this end, we established intrapulmonary ABN delivery as a means of eliminating activation in blood and off-target organs (reducing noise), while maximizing delivery to the target organ (increasing signal) (Fig. 1A-B). After cleavage of ABN substrates by proteases in the lung, reporters rapidly entered the urine via the blood, where they were quantified by mass spectrometry (Fig. 1C-D). Finally, we leveraged a machine learning classification algorithm, termed random forest, to achieve diagnostic sensitivity of 92% and specificity of 100% in detecting early-stage disease in a genetically engineered, *Kras* and *Trp53* mutant mouse model of lung adenocarcinoma (Fig. 1E).

**Fig. 1.**
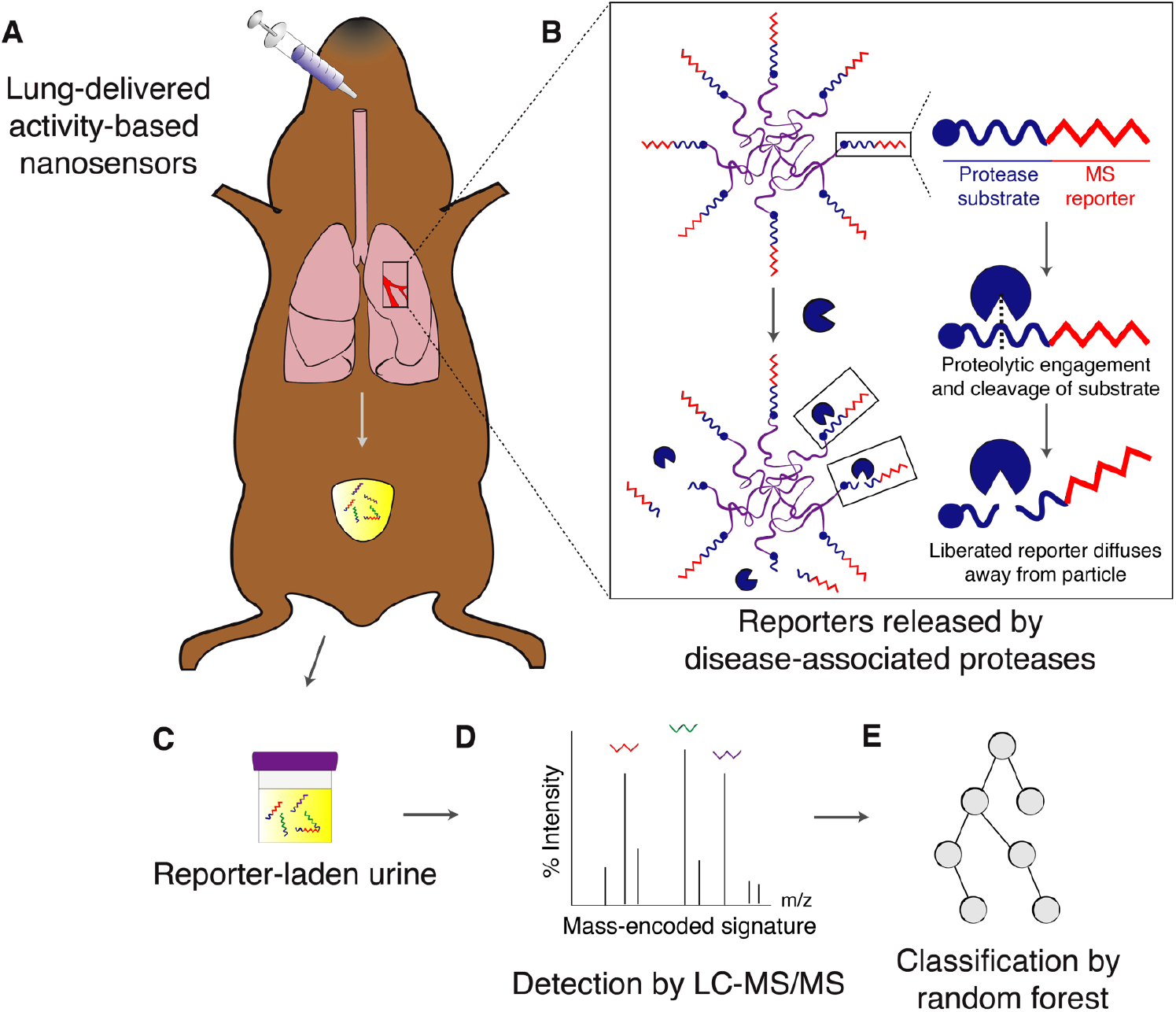
Approach and overview. (**A**) ABNs are administered intratracheally and reach the lung epithelium. (**B**) At the tumor periphery, disease-associated proteases cleave protease substrates, liberating mass-encoded (MS) reporters from the PEG scaffold. (**C**) These reporters are small enough to diffuse into the bloodstream and passively filter into the urine for detection. (**D**) Synthetic disease reporters are detected in the urine by liquid chromatography followed by tandem mass spectrometry (LC-MS/MS). (**E**) Random forest classification is performed on a training cohort of mice and subsequently tested on an independent validation cohort in order to provide a positive or negative diagnosis of malignancy.

## Results

### *Proteases are overexpressed in a* Kras *and* Trp53 *mutant mouse model of lung adenocarcinoma*

Common driver mutations of NSCLC in humans are those that activate *KRAS* (10-30%) or inactivate function of *TP53* (50-70%)^32^. To examine the ability of ABNs to detect lung cancer in a relevant mouse model, we selected a genetically driven model of adenocarcinoma (a type of NSCLC that accounts for 38% of all cases of lung cancer^33^) that incorporates mutations in these genes. This extensively characterized model uses intratracheal administration of virus expressing Cre recombinase to activate mutant *Kras^G12D^* and delete both copies of *Trp53* in the lungs of *Kras^LSL-G12D/+^;Trp53^fl/fl^* (KP) mice (fig. S1A), initiating tumors that closely recapitulate human disease progression from alveolar adenomatous hyperplasia to grade IV adenocarcinoma over the course of about 18-20 weeks (fig. S1B)^34^.

In anticipation of our use of the KP model to validate ABNs *in vivo*, we sought to characterize protease expression in tumor-bearing KP mice to nominate protease targets. To that end, we selected a recently published RNA-Seq dataset that profiled KP tumors across disease stages, and we used it to identify overexpressed secreted protease genes^35^. In this study, tumor cells expressing a fluorescent reporter had been isolated by FACS and profiled by RNA-Seq. We pooled samples from metastatic (T_met_, *n* = 9), non-metastatic (T_non-met_, *n* = 10), and early stage (KP-Early, *n* = 3) tumors, as well as Kras-mutant, Trp53-intact (K, *n* = 3) tumors and identified proteases that were overexpressed in tumor cells relative to normal lung cells (*n* = 2) (Fig. 2A).

**Fig. 2.**
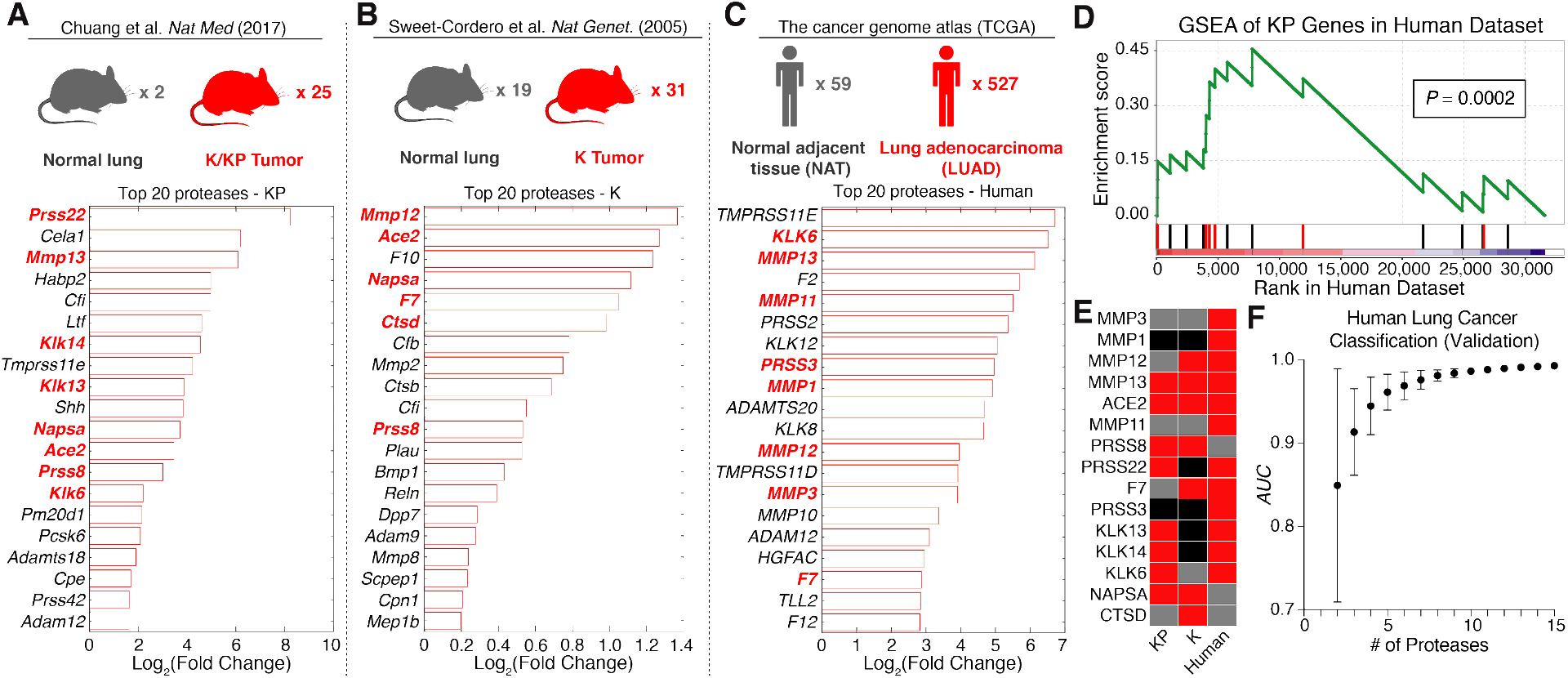
Proteases are overexpressed in lung cancer and enable classification of human disease. (**A-C**) Existing RNA-Seq (A,C) and microarray (B) datasets were analyzed to identify extracellular endoproteases overexpressed in human and murine lung cancer. Gene expression fold changes in lung cancer compared to control lung tissue were calculated by FPKM in the KP dataset (A), significance analysis of microarrays (SAM) in the K dataset (B), and DESeq2 in the human dataset (C). Protease genes in red are those that were selected for the “LUAD protease panel”. (**D**) Gene set enrichment analysis (GSEA) was performed in the TCGA (human) dataset using the top 20 overexpressed protease genes in KP tumors. Red bars are genes included in the “LUAD protease panel”. The maximum enrichment score was 0.455 (*P* = 0.0002). (**E**) A set of 15 proteases was selected as the “LUAD protease panel”. Red: Fold_Disease_ > 1· Grey: Fold_Disease_ < 1. Black: Not included in dataset. (**F**) Generalized linear model classification was performed in the TCGA dataset using the 15 protease genes in the “LUAD protease panel” as features. Area under the receiver operating characteristic curve (AUC) for the validation cohort is shown as a function of the number of proteases included in the classifier (*n* = 50 combinations of protease genes for each point). Error bars represent SD.

Because this dataset was derived from FACS-purified tumor cells, it failed to take into account contributions from the KP tumor microenvironment. In addition, it was limited in its representation of early-stage disease. We therefore analyzed an additional gene expression dataset profiling the K model^36^, which is transcriptionally similar to early-stage KP tumors and human lung adenomas^35^. Significance analysis of microarrays (SAM) was used to identify proteases with increased expression in K tumors relative to normal lungs^37^ (Fig. 2B).

### Proteases overexpressed in the KP mouse model are relevant to human lung adenocarcinoma

To ensure that ABNs were tuned to address human lung adenocarcinoma (LUAD)-associated proteases^38–40^ in addition to proteases enriched in the KP model, we mined The Cancer Genome Atlas (TCGA) dataset, with mRNA sequencing (RNA-Seq) and clinical data collected from 527 LUAD patients^41^. We analyzed expression levels of 168 candidate human extracellular endoprotease genes in these patients using the DESeq2 differential expression analysis package (Fig. 2C)^42^. Of the 527 TCGA patients with RNA-Seq data for primary LUAD (294 stage I, 123 stage II, 84 stage III, and 26 stage IV), 59 had matched normal adjacent tissue suitable for use as a comparison (Fig. 2C, top). Of the 20 most highly upregulated proteases, nine were metalloproteases, 11 were serine proteases, and several overlapped with proteases overexpressed in KP tumors (Fig. 2C, bottom).

We then sought to assess whether proteases associated with benign lung diseases could confound the specificity of ABNs for lung cancer. To this end, we performed receiver operating characteristic (ROC) analysis on RNA-Seq data from interstitial lung disease (ILD) and chronic obstructive pulmonary disease (COPD), curated by the Lung Genomics Research Consortium (LGRC). In ROC analysis, the sensitivity and specificity of a given classifier (e.g. protease gene expression) in discriminating between two cohorts (e.g. disease and control) are assessed across a series of cutoff values. The area under the curve (AUC) is then calculated as measure of classification accuracy, where a perfect diagnostic has an AUC of 1 and a random diagnostic has an AUC of 0.5. ROC analysis revealed that proteases overexpressed in LUAD were not increased in COPD or ILD (fig. S2A)^43^; none of the 10 proteases included in the analysis classified benign lung diseases from healthy lungs with an AUC greater than 0.6. In contrast, classification efficiency in LUAD reached above 0.9 in eight out of ten cases (fig. S2B-D). The finding that genes upregulated in LUAD are not overexpressed in COPD or ILD may be due to our use of NAT as “normal” tissue when nominating proteases for the panel, as NAT is known to harbor inflammatory gene expression changes that distinguish it from “true normal” tissue^44^. Therefore, the genes of the LUAD protease panel are more likely to be specific to cancer, rather than inflammation or other nonspecific disease-associated processes.

To assess whether the proteolytic landscape of the KP model recapitulates that of human lung cancer, we performed gene set enrichment analysis (GSEA)^45^ in the TCGA dataset using the top 20 overexpressed proteases in the KP model (Fig. 2D). GSEA assesses the extent to which a particular gene set (S) is enriched in a gene expression dataset by rank-ordering all genes in the dataset and iterating through the list, increasing the enrichment score each time a gene in *S* is encountered, and decreasing it otherwise. This approach revealed significant enrichment of 14 of the top 20 KP-expressed protease genes in human lung adenocarcinoma, yielding a maximum enrichment score of 0.455 (*P* = 0.0002).

### A panel of proteases overexpressed in human and mouse lung adenocarcinoma enables robust classification of human disease

A set of 15 proteases overexpressed across all or a subset of the mouse and human datasets was then selected as a “LUAD protease panel”, consisting of six metallo-, seven serine, and two aspartic proteases (Fig. 2E, and indicated in bold red text in Fig. 2A-C). We next returned to the TCGA dataset and evaluated the performance of this panel in classifying human lung cancer from healthy lungs on the transcriptional level. Generalized linear model classification was performed using the Caret package, using the 15 LUAD proteases as features. ROC analysis revealed that the AUC increases with increasing information (i.e. number of proteases), achieving nearly perfect classification in the validation cohort with all 15 proteases (Fig. 2F).

### Cleavage of multiplexed substrate panel follows class-specific patterns

We have previously designed and validated hundreds of peptide sequences as protease substrates, leveraging known catalytic specificities of different protease families, published datasets, and substrate sequences in databases like cutDB and MEROPS^46,47^. We nominated 14 of these substrates in an effort to encompass the cleavage preferences of metalloproteases (MP), serine proteases (SP), and aspartic proteases (AP), all of which were included in our LUAD protease panel, and characterized the catalytic reactivity of each protease-substrate pair. We synthesized quenched probes that incorporated the 14 peptide substrates (PPQ1-14), such that they fluoresce upon proteolytic cleavage to enable real-time monitoring of protease activity *in vitro* (Fig. 3A and table S1). We incubated each individual probe with each protease in the LUAD panel and measured protease activity by monitoring fluorescence increase over the course of 45 minutes. Shown are sample kinetic plots monitoring proteolytic dequenching of the 14 FRET-paired probes when incubated with (above) and without (below) recombinant matrix metalloprotease 3 (MMP3) (Fig. 3B). We found that hierarchical clustering of fluorescence fold changes of each substrate led to the separation of proteases of different classes (Fig. 3C). We also found that while certain probes were cleaved selectively by individual classes of protease (e.g. PPQ2 and PPQ11 for MP and SP, respectively), others were cleaved well by proteases of multiple classes (e.g. PPQ3, PPQ12 for MP/AP and MP/SP, respectively) (fig. S3). Overall, the dequenching panel results indicated that the set of 14 probes provided robust coverage of the cleavage profiles of all three classes represented by the LUAD protease panel.

**Fig. 3:**
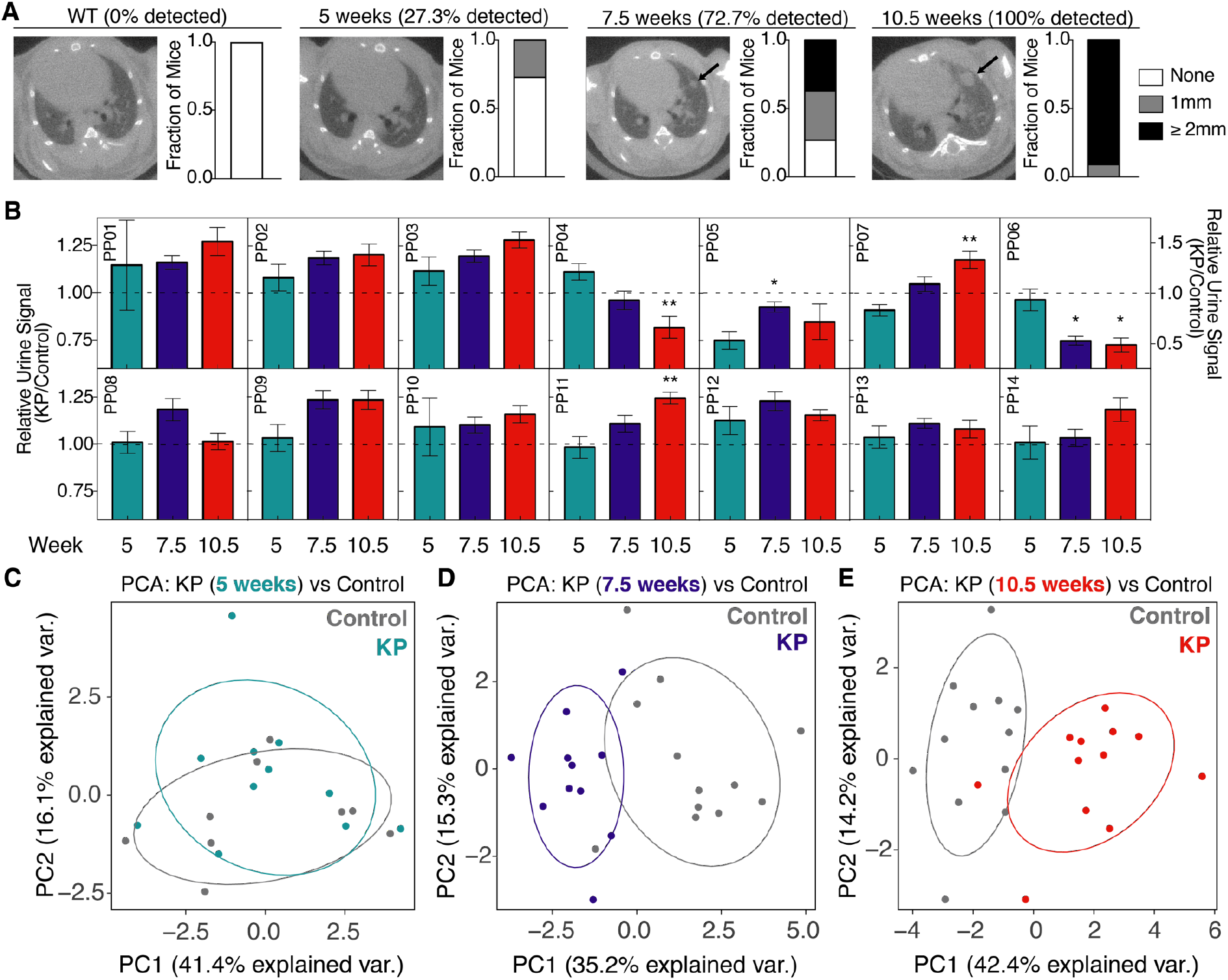
LUAD substrate panel cleavage patterns are driven by protease class. (**A**) All 15 proteases in the “LUAD protease panel” were screened against a panel of 14 FRET-paired (quenched) protease substrates and fluorescence activation was monitored over 45 minutes. (**B**) Kinetic fluorescence curves are shown for 14 FRET-paired substrates with (upper panel) and without (lower panel) addition of MMP3. (**C**) Fluorescence fold changes at 45 minutes (average of 2 replicates) were tabulated and hierarchical clustering was performed to cluster proteases (vertical) by their substrate specificities and substrates (horizontal) by their protease specificities. Proteases labeled in green, orange or blue represent metallo, serine or aspartic proteases, respectively.

### Pulmonary-delivered nanoparticles distribute throughout the lung and reach the tumor periphery

The lung efficiently and rapidly exchanges compounds with the bloodstream owing to high surface area and very thin barriers; human adult lungs have an area of ~100 m^2^ and, in alveoli, type I cells can be <0.1 μm thick^48^. Inhaled molecules and particles cross into the bloodstream by passive diffusion, transcytosis, or paracytosis, with rate and route of transit largely dependent on size and hydrophobicity^48^.

To adapt the ABN platform for highly sensitive and specific detection of early-stage lung cancer, we sought to circumvent background protease activity present in the blood and off-target organs, which can nonspecifically liberate reporters, by administering the nanosensors via localized intrapulmonary, rather than systemic intravenous, delivery. We built ABNs using a 40 kDa eight-arm poly(ethylene glycol) (PEG-8_40kDa_) nanoparticle coupled to protease substrates bearing terminal mass-encoded reporters (Fig. 1B). Similarly sized PEG particles have been shown to remain in the lung with half-lives of several hours and relatively little phagocytosis^49^; consequently, we anticipate that the ABNs are largely free to sample extracellular lung protease activity over the time period during which we monitor urinary reporter accumulation. To assess biodistribution of ABNs following intrapulmonary delivery, we labeled the PEG-8_40kDa_ scaffold with a near-infrared dye, VivoTag750, delivered the nanoparticles to mice by intratracheal (IT) intubation or intravenous (IV) injection, and collected organs after 60 minutes (Fig. 4A). Fluorescence imaging revealed deep delivery of nanoparticles to all lung lobes in mice receiving IT particles, but negligible delivery to other organs (Fig. 4B-C). In contrast, only 14% of organ fluorescence was confined to the lung in the IV-delivered group. In terms of absolute delivery of ABNs, lung fluorescence in the IT group was 263 times greater than liver fluorescence (*P* < 0.0001), while lung fluorescence was 30% lower than liver fluorescence in the IV group. As blood is a rich, non-specific proteolytic matrix and achieving organ-specific biodistribution of systemically delivered nanoparticles remains difficult, IT ABNs offer distinct advantages over IV-delivered variants.

**Fig. 4:**
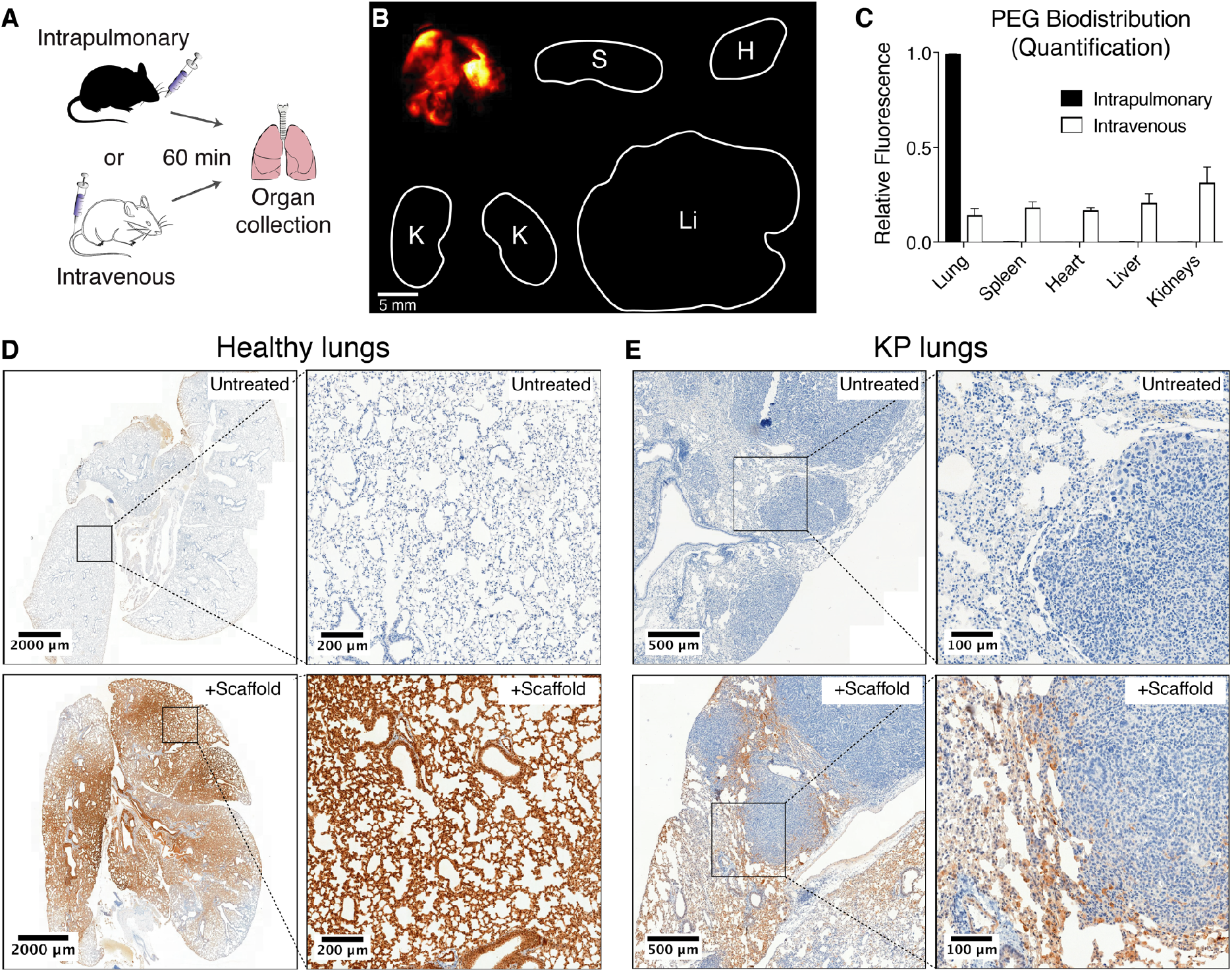
Intrapulmonary-administered nanoparticle scaffolds penetrate deep within the lung and reach the periphery of KP tumors. (**A**) Wild-type mice were treated intratracheally (IT) or intravenously with VT750-labeled PEG-8_40_kDa and biodistribution was assessed. (**B**) Fluorescent imaging of organs was performed 60 min post-IT delivery. Clockwise from top-left: lung, spleen, heart, liver, kidneys. (**C**) Organ-specific biodistribution was quantified (*n* = 4 each condition). Error bars represent SD. (**D**) Healthy mice were either untreated (above, *n* = 1) or treated with IT administration of biotin-labeled PEG scaffold (below, *n* = 2), followed by excision of lungs and immunohistochemical staining for biotin (brown). (**E**) Advanced-stage (16.5 week) KP mice were either untreated (top, *n* = 3) or treated with IT administration of biotin-labeled PEG scaffold (bottom, *n* = 3), followed by excision of lungs and immunohistochemical staining as in (D).

To assess microscopic distribution of the ABN scaffold within the lung following IT delivery, we labeled the PEG-8_40kDa_ scaffold with biotin and administered the nanoparticles to healthy mice by intratracheal intubation. Lungs were collected from mice 20-30 minutes post-IT delivery, fixed, and stained for biotin. While lungs from untreated mice were negative for biotin (Fig. 4D, top), lungs from mice that received the scaffold demonstrated broad distribution of nanoparticles throughout the lung (Fig. 4D, bottom left) and specifically within terminal alveoli (Fig. 4D, bottom right).

We then administered biotin-labeled PEG-8_40kDa_ scaffold in high grade KP tumor-bearing mice, by intratracheal intubation, to assess whether these particles are able to reach the site of disease. Again, while lungs from untreated KP mice were negative for biotin (Fig. 4E, top), lungs from KP mice that received intrapulmonary delivery of the biotinylated scaffold demonstrated presence of nanoparticles at the margin of tumors where protease activity is relevant to disease growth and invasion^28–30^ (Fig. 4E, bottom).

### Mass-encoded reporters filter from the lung to the urine via the blood and are detectable by mass spectrometry

In order to enable multiplexed detection of a broad spectrum of disease-associated proteases via a single *in vivo* administration of nanosensors, we conjugated each member of the LUAD substrate panel to a uniquely identifiable mass-encoded reporter (PP1-14; Table 1). Following substrate proteolysis, the encoded reporters diffuse away from the nanoparticle scaffold and, due to their small size, efficiently cross into the bloodstream and are subsequently concentrated into the urine by glomerular filtration (Fig. 1B). As previously described^19^, we used variable labeling of the 14-mer Glu-Fibrinopeptide B (Glu-Fib) with stable isotope-labeled amino acids to uniquely barcode each of the 14 peptide substrates. Multiple reaction monitoring via a liquid chromatography triple quadrupole mass spectrometer (LC-MS/MS) enables quantitative assessment of peptide-liberated urinary reporter concentration within a broad linear range (1-1000 ng/mL, fig. S4A). To assess the efficiency of urinary accumulation of reporters after liberation from the PEG scaffold (termed “free reporters”), we administered mass-encoded free reporters by IT and IV administration, collected urine, and performed LC-MS/MS. We found that urinary accumulation scaled linearly with input doses between 2.5 ng and 25 ng for both routes of delivery (slope_IT_ = 0.075 ng^−1^, slope_IV_ = 0.077 ng^−1^; fig. S4B). We also investigated pharmacokinetics of the free reporter by administering a Cy7-labeled version of Glu-Fib (the cleavage product after liberation from the PEG scaffold) both IT and IV. Pharmacokinetic data revealed characteristic single-exponential concentration decay following intravenous injection (fig. S4C). In contrast, the pharmacokinetic behavior of the free reporter following IT administration is suggestive of an initial phase of partitioning from the alveoli into the blood (peaking at 1 to 2 hours after delivery), followed by renal filtration from the blood.

**Table 1.**
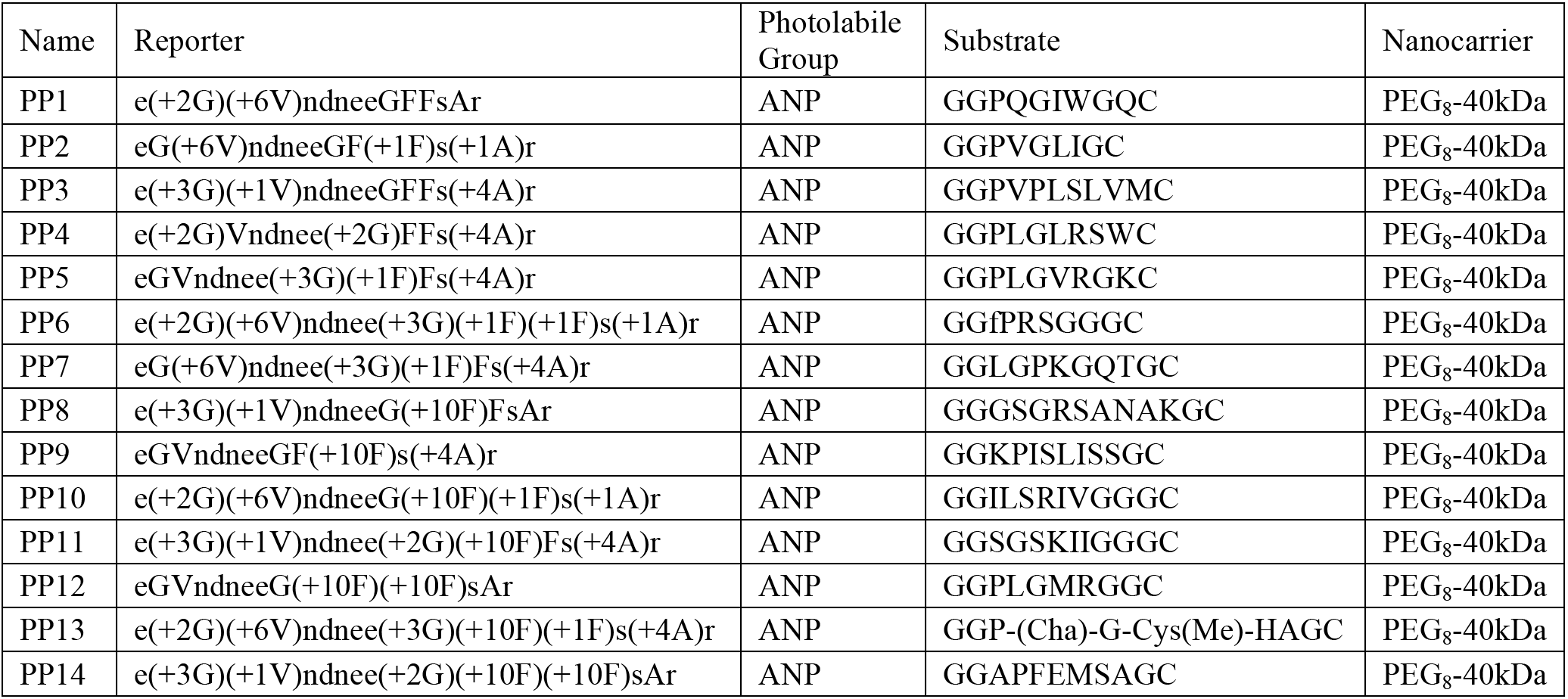
Reporter and substrate sequences for *in vivo* urinary diagnostics. ANP, 3-Amino-3-(2-nitro-phenyl)propionic Acid; Cha, 3-Cyclohexylalanine; Cys(Me), (methylsulfanyl)propanoic acid; lowercase letters, D-amino acids

### Early-stage lung tumors in the KP model are detectable by ABNs

With the observation that IT delivery of mass-encoded reporters leads to their partitioning from lung to circulation and subsequent concentration in the urine, we sought to longitudinally monitor disease progression in KP mice with ABNs and benchmark their diagnostic performance against microCT. After initiating disease via administration of adenovirus, we monitored development of tumor burden by performing microCT at 5 weeks, 7.5 weeks, and 10.5 weeks (Fig. 5A, representative microCT slice at each time point, with arrow indicating development of a single nodule over time). At 5 weeks, only grade 1 tumors are present in the KP model, while at 7.5 and 10.5 weeks, grade 2 disease is expected (fig. S1B)^34^. Tumor burden was quantified on microCT by a blinded radiation oncologist at each time point (maximum nodule size shown in bar graph form to the right of each image). Median nodule multiplicity by microCT was 0 (range 0-3) at 5 weeks, 2 (range 0-6) at 7.5 weeks, and 4 (range 1-8) at 10.5 weeks. The sensitivity of microCT at 100% specificity was 27.3% at 5 weeks, 72.7% at 7.5 weeks, and 100% at 10.5 weeks.

**Fig. 5:**
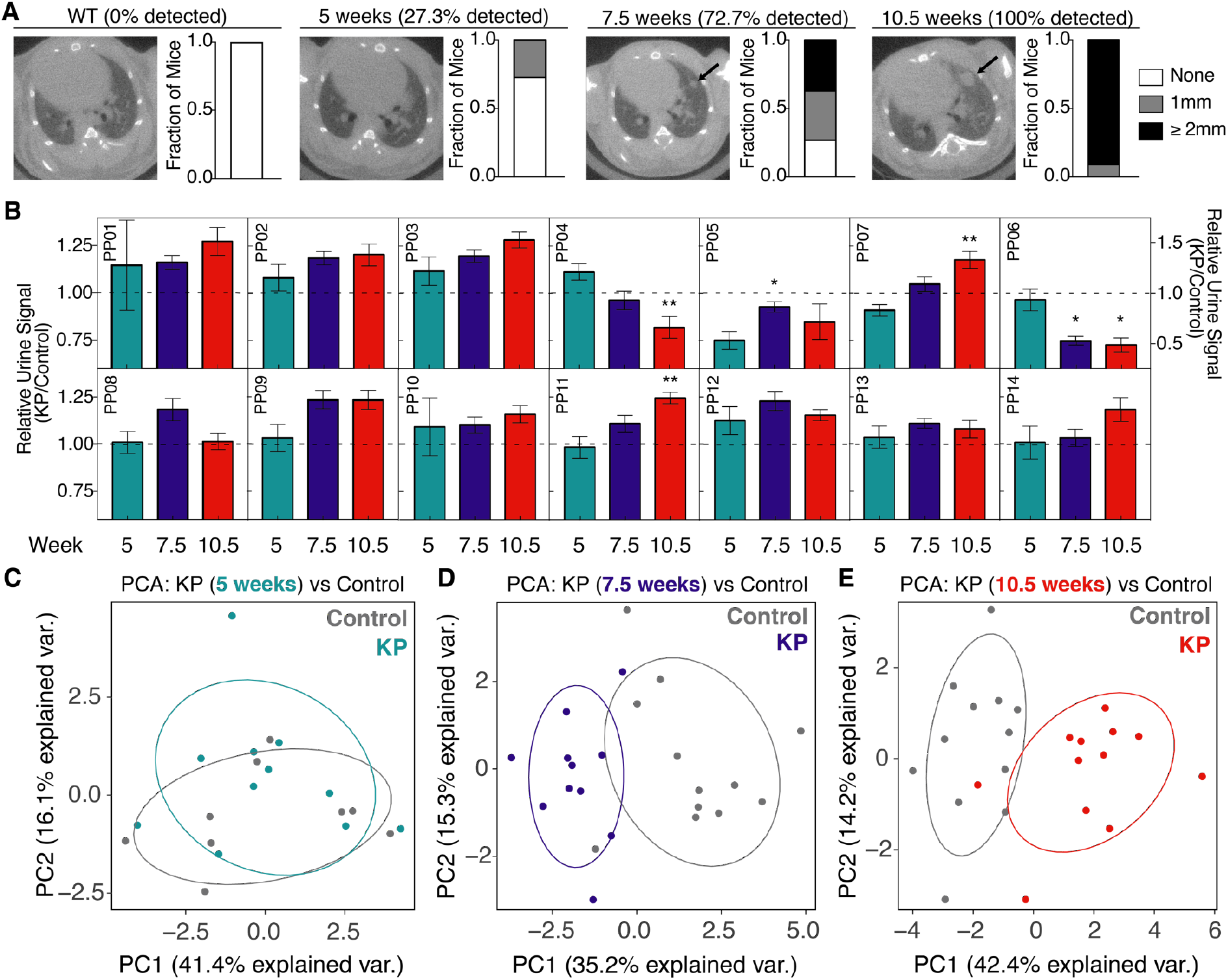
ABNs distinguish between diseased and healthy mice. (**A**) Tumor development was monitored by microCT in healthy (left, *n* = 11) and KP mice at 5 weeks (*n* = 11), 7.5 weeks (*n* = 11), and 10.5 weeks (n = 11) after tumor induction. Right three panels represent time series of a single mouse, with arrow indicating development of a single nodule over time. Size of the largest tumor nodule was assessed by a blinded radiation oncologist (quantification at right of each image). (**B**) ABNs were administered to KP and control animals at 5 weeks (KP: *n* = 11; Control: *n* = 9), 7.5 weeks (KP: *n* = 11; Control: *n* = 12), and 10.5 weeks (KP: *n* = 12; Control: *n* = 12) after tumor initiation, bladder was voided at 1 hr, and urine was collected and pooled over the following 1 hour interval. LC-MS/MS was performed, peak area ratio (PAR, peak area of reporter divided by peak area of spiked-in internal standard) was calculated, and all reporters were mean normalized within each sample. Y axis represents 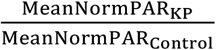 for each reporter at each time point. For clarity, PP06 is presented on a larger scale y axis. Asterisks indicate significant differences from 5 weeks. * *P_adj_* < 0.05, ** *P_adj_* < 0.01; by two-tailed *t*-test with adjustment for multiple hypotheses using the Holm-Sidak method. Error bars represent SEM. (**C-E**) Unsupervised clustering by principal component analysis (PCA) was performed on mean normalized MS data for KP mice and controls at 5 weeks (C), 7.5 weeks (D), and 10.5 weeks (E).

To characterize ABN performance relative to microCT *in vivo*, we administered all 14 protease-sensitive ABNs to the lungs of KP mice and healthy, age and sex-matched controls at 5, 7.5, and 10.5 weeks after tumor initiation. Mouse bladders were voided one hour after intrapulmonary delivery and all fresh urine produced during the subsequent hour (from 60-120 minutes after ABN administration) was pooled and collected. LC-MS/MS was performed and peak area ratios (defined as peak area of urinary reporter divided by peak area of spiked-in internal standard) of protease-sensitive reporters were mean-normalized within each urine sample to reduce mouse-to-mouse variation. Several reporters differentiated KP mice from the healthy control group, with some reporter differences becoming amplified over time (e.g. PP07, PP11) (Fig. 5B). At 7.5 weeks and 10.5 weeks, 5/14 reporters were significantly different between KP and healthy mice (*P_adj_* < 0.05), while none of the reporters differed at 5 weeks (fig. S5). Three of the 5 reporters enriched in KP urine were the same at 7.5 and 10.5 weeks (PP02, PP03, and PP09), and these corresponded to peptides cleaved by metallo or both metallo and aspartic proteases *in vitro*. However, the most significantly enriched reporter in the urine of KP mice at 10.5 weeks (PP11) corresponded to a peptide cleaved only by serine proteases *in vitro*. Unsupervised clustering by principal component analysis (PCA) succeeded in separating most KP and control mice at the 7.5 week and 10.5 week time points, but not at 5 weeks (Fig. 5C-E).

### Machine learning classification enables sensitive and specific disease diagnosis

As a step toward clinical translation of ABNs as a prospective diagnostic tool, we sought to demonstrate that a classifier could be trained on a subset of healthy and tumor-bearing mice and validated on an independent cohort. We trained a random forest classifier using the ABN reporter output from 50% of control mice tested at 5 weeks, 7.5 weeks, and 10.5 weeks, as well as from 50% of the KP mice tested at 7.5 weeks (Fig. 6A). Random forest is a high-performance classifier, applicable to a wide variety of classification tasks, that generates a collection of decision trees (a “forest”) that are sampled to produce classification results^50^. The classifier assigned a probability that each mouse belonged to either the KP cohort or the healthy control cohort (Fig. 6B, “Training” panel), achieving perfect separation of control and KP mice. We then locked and tested the classifier on an independent validation cohort consisting of classifier-naïve KP mice assayed at 5 weeks, 7.5 weeks, and 10.5 weeks post-induction, as well as the remaining control mice from each time point. KP mice were significantly more likely to be classified as “KP” than were control mice at 7.5 weeks and 10.5 weeks but not at 5 weeks (Fig. 6B, “Validation” panel). Accordingly, ROC analysis on the validation subset of this probability data revealed no classification power at 5 weeks (*AUC_5wks_* = 0.58, *P* = 0.7) but significant classification at 7.5 weeks and 10.5 weeks (*AUC_7.5wks_* = 0.96, *P* = 0.02; *AUC_10.5wks_* = 0.95, *P* = 0.0005) (Fig. 6C). With 100% specificity, ABNs exhibited sensitivity of 80% at 7.5 weeks and 92% at 10.5 weeks, outperforming microCT in the detection of millimeter-scale tumors at 7.5 weeks (Fig. 5A). Together, these data illustrate the power of multiplexed, lung-specific ABNs to intercept lung tumors early in disease development.

**Fig. 6.**
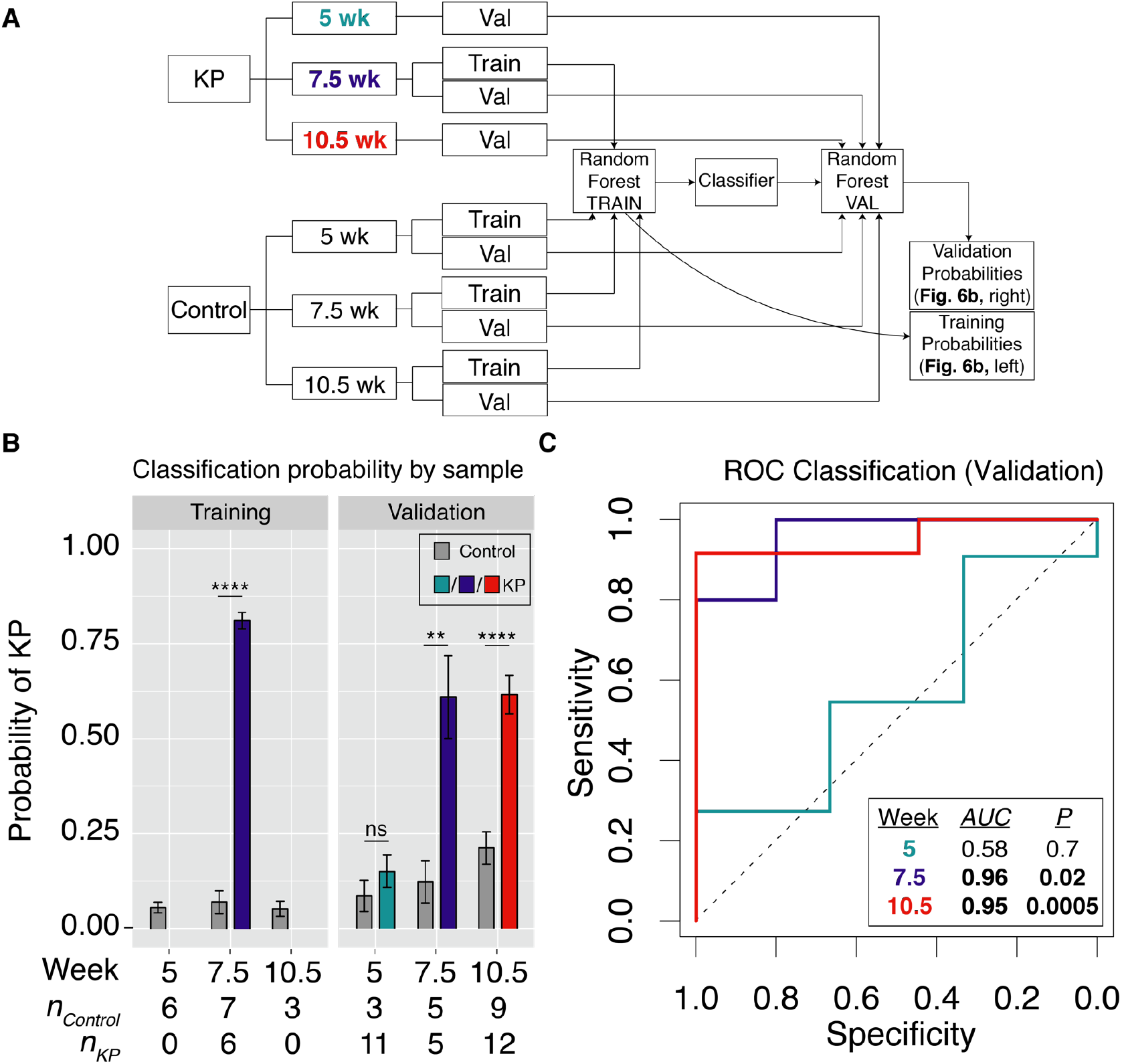
ABNs enable highly sensitive and specific detection of early-stage lung cancer. (**A**) Schematic of approach. Random forest classifier was trained on mean normalized urinary reporters from KP mice at 7.5 weeks, as well as control mice at 5 weeks, 7.5 weeks, and 10.5 weeks. Classifier was validated on KP mice and control mice at all 3 time points. (**B**) Random forest classifier returned the probability that each mouse was either “Control” or “KP”. ** *P* < 0.01, **** *P* < 0.0001; by two tailed *t*-test. Error bars represent SEM. (**C**) ROC analysis was performed on probability data to generate AUC values for the validation cohorts at 5 weeks (*AUC* = 0.58), 7.5 weeks (*AUC* = 0.96), and 10.5 weeks (*AUC* = 0.95).

## Discussion

In this work, we present an advance toward clinical translation of a new class of biomarkers, ABNs. We found that multiplexed ABNs, when delivered by intratracheal instillation, performed with diagnostic sensitivity of up to 92% and specificity of 100% for local, early-stage disease in an immunocompetent, genetically engineered, *Kras* and *Trp53* mutant lung adenocarcinoma model (Fig. 6C). Importantly, this model recapitulates the proteolytic landscape of human lung cancer (Fig. 2D) and is notable for overexpression of key enzymes associated with human disease, including MMP13 and several kallikreins (Fig. 2A, C). Our approach overcomes the intrinsic sensitivity limitation of blood-based diagnostic assays for early-stage disease by profiling disease activity directly within the tumor microenvironment and providing multiple steps of signal amplification^31^. We further ensure, by delivery via intratracheal instillation, that virtually all ABNs reach the lung and bypass nonspecific activation in off-target organs (Fig. 4B-C).

Improved diagnostic tools are needed for lung cancer, as most patients present to the clinic when their disease has reached too advanced a stage for potentially curative therapy (e.g. surgical resection and/or chemoradiation) to be administered^1^. Screening by LDCT results in high false positive rates, leading many healthy patients to undergo unnecessary follow-up procedures that are costly and invasive^4^. Though there exist no widely accepted endogenous biomarkers for lung cancer^51^, several recently reported molecular diagnostic strategies hold promise. Multiplexed ctDNA profiling may be combined with protein biomarkers (as in CancerSEEK) to improve diagnostic accuracy for early-stage disease^52^. However, while the sensitivity of this approach is high for certain cancer types, it is modest at 60% for lung cancer. Analysis of volatiles in exhaled breath (by mass spectrometry or nanosensor arrays) has been shown to distinguish lung cancer patients from healthy controls, but further validation is needed to verify the specificity of these volatiles for malignant, rather than benign, pulmonary diseases^53,54^. Combining ABNs with orthogonal diagnostic approaches that leverage ctDNA, protein biomarkers, CTCs, and/or volatiles may enhance diagnostic accuracy over any one modality alone.

This study represents a significant step toward clinical implementation of ABNs, validating the efficacy of the tool in an early-stage, immunocompetent GEMM, rather than a flank xenograft model. Though a preclinical model cannot fully recapitulate the heterogeneity of human lung cancer, a GEMM offers several advantages over a cell transplant model. In addition to its genetic and phenotypic homology to human lung cancer, the KP model enables evaluation of immune cell-associated protease activity, as well as assessment of stage-specific differences in proteolytic signatures. For instance, though metalloprotease-sensitive ABNs are, as expected, preferentially cleaved in KP lungs at both 7.5 and 10.5 weeks, the activation of PP11 (a serine protease-sensitive substrate, fig. S3) in KP mice 10.5 weeks after disease induction could point to an unexpected role of serine protease activity in tumor progression at this disease stage (fig. S5). An intriguing hypothesis is that endogenous serine protease inhibitors may be downregulated during the maturation of primary tumors. Indeed, maspin is a serine protease inhibitor known to inhibit the metastatic and angiogenic potential of tumor cells^55,56^ and its transcript (*Serpinb5*) is significantly downregulated in pleural, soft tissue, and liver metastases relative to primary tumors in the KP model^35^. Dysregulated coagulation, which is driven by a cascade of serine proteases, has also been implicated in tumorigenesis and could drive ABN activation *in vivo^57^*. Transcriptomic, proteomic, and *ex vivo* protease activity assays could be leveraged to rigorously define the mechanistic underpinnings of such stage-specific differences in ABN cleavage patterns.

In this work, we have demonstrated the sensitivity of intrapulmonary ABNs for local, early-stage lung cancer, but a challenge to overcome prior to clinical implementation is ensuring the specificity of ABNs for cancer over benign lung diseases. Though we provide preliminary evidence that proteases associated with lung cancer are not overexpressed in COPD or ILD on the RNA level (fig. S2), further validation in mouse models and human samples will be needed. We can further improve the specificity of ABNs for malignancy by screening a large, diverse panel of peptide substrates, via high throughput methods like substrate phage^58^ or CLiPS^59^ display, against *ex vivo* biospecimens from patients with LUAD, COPD, ILD, granulomas, and hamartomas^60^. Substrates can then be downselected on the basis of preferential cleavage by LUAD tissue to yield a highly specific panel. In parallel, we will also explore the compatibility of ABNs with pulmonary delivery systems like dry powder inhalers and nebulizers, with an eye toward clinical implementation.

In summary, intrapulmonary ABNs perform with high sensitivity and specificity for detection of local, early-stage lung cancer in a GEMM, via a non-invasive urine test. This performance is enabled by integrating across gene expression datasets of human and mouse lung adenocarcinoma to identify candidate proteases, screening these candidates against FRET-paired peptide substrates *in vitro*, and directly delivering ABNs incorporating these substrates into the lungs of mice. Future efforts will center on exploring protease biology at the earliest stages of lung cancer development in humans, designing ABNs that are highly specific for these proteases via high throughput screening methods, and evaluating their responsiveness in human biospecimens. Clinically, ABNs may be used in conjunction with LDCT to enhance specificity and reduce the number of patients referred for invasive follow up procedures. With further optimization and validation studies, ABNs may one day provide an accurate, noninvasive, and radiation-free strategy for screening.

## Materials and Methods

### Study design

The goal of this study was to determine whether intrapulmonary administration of a multiplexed library of ABNs could be used to detect localized, early-stage, lung cancer. All mouse studies were approved by the MIT committee on animal care (protocol 0414-022-17) and were conducted in compliance with all MIT ethical policies. Experiments involving intrapulmonary delivery of ABNs in KP mice consisted of 12 KP mice and 12 healthy control mice, and these mice were monitored, by intratracheal ABN administration and microCT, at 5 weeks, 7.5 weeks, and 10.5 weeks after tumor induction. Sample size was selected to ensure a sample size greater than or equal to five for both training and validation groups at each time point and for each treatment group. Urine samples with peak area ratio (PAR) values of zero for two or more analytes were excluded, as these samples represented failed ABN deliveries and would confound analysis. For differential expression analysis of protease genes in KP mice, genes for which neither normal lung sample was nonzero were excluded, as calculation of fold changes (Tumor/Normal) would otherwise yield undefined values. For AUROC analysis in the LGRC dataset, genes for which greater than half of the samples had FPKM values of zero were excluded. During selection of KP and healthy control mice from the Jacks Lab breeding colony, we were blinded to all characteristics but age, sex, and genotype. For random forest classification, mice were randomly assigned to the training and validation cohorts using a randomly generated seed.

### Gene expression analysis

Human RNA-Seq data was generated by the TCGA Research Network (http://cancergenome.nih.gov; all 527 primary lung adenocarcinoma cases^41^) and the Lung Genomics Research Consortium (LGRC; all 89 patients^43^). The list of human extracellular protease genes was obtained from UniProt using the following query: (keyword:“Protease [KW-0645]”) locations:(location:secreted) AND reviewed:yes AND organism:“Homo sapiens (Human) [9606]”. Differential expression analysis on the TCGA data was performed using the DESeq2 differential expression library in the R statistical environment^42,61^. Area under the receiver operating characteristic curve (AUROC) analysis was performed for the TCGA and LGRC datasets using FPKM values from disease samples (LUAD, ILD, and COPD) and their respective controls (NAT for LUAD, normal lung for ILD and COPD), using GraphPad Prism version 7.0a for Mac OS X, GraphPad Software, La Jolla California USA, www.graphpad.com. Genes in the LGRC dataset for which at least half of the samples had FPKM values greater than zero were included in the AUROC analysis, but all zero values were excluded. FPKM values for the KP model^35^ were downloaded from GEO. Top 20 extracellular endoproteases were identified by averaging FPKM values across all tumor bearing mice (K, KP-Early, T_non-met_, and T_met_) and dividing by the average FPKM values for normal mice. Genes for which neither of the two normal lung samples had nonzero FPKM values were excluded. Microarray counts for the K dataset^36^ were downloaded from GEO. Gene expression fold changes were determined by performing quantitative significance analysis of microarrays (SAM) using the “Standard” regression method, 100 permutations, and 10 neighbors for KNN^37^.

Pre-ranked gene set enrichment analysis (GSEA) was performed on the LUAD gene expression dataset from TCGA, using a gene set containing the top 20 overexpressed proteases in the KP model^35^. The pre-ranked list of log_2_(Fold Change) was generated previously by DESeq2. 10000 permutations by gene set were performed to calculate the *P* value. GSEA was performed via the GenePattern online software^62^.

### Fluorogenic substrate characterization

Fluorogenic protease substrates were synthesized by CPC Scientific. Recombinant proteases were purchased from Enzo Life Sciences, R&D Systems, and Haematologic Technologies. For recombinant protease assays, fluorogenic substrates PPQ1-14 (1 μM final concentration) were incubated in 30 μL final volume in appropriate enzyme buffer, according to manufacturer specifications, with 12.5 nM recombinant enzyme at 37°C. Proteolytic cleavage of substrates was quantified by increases in fluorescence over time by fluorimeter (Tecan Infinite M200 Pro). Enzyme cleavage rates were quantified as relative fluorescence increase over time normalized to fluorescence before addition of protease. Hierarchical clustering was performed in GENE-E, using fluorescence fold changes at 45 minutes.

### Biodistribution studies

For all mouse experiments, anesthesia was induced by isofluorane inhalation (Zoetis) and mice were monitored during recovery. Biodistribution studies were performed in C57BL/6 mice. VT750-NHS Ester (PerkinElmer) was coupled to 8-arm 40 kDa PEG-amine (PEG-8_40kDa_-amine, JenKem) at a 4:1 molar ratio, reacted overnight, and purified by spin filtration. Mice were lightly anesthetized via isoflurane inhalation and PEG-8_40kDa_-VT750 (50 uL volume, 5 uM concentration by VT750 absorbance) was administered by passive inhalation following intratracheal intubation with a 22G flexible plastic catheter (Exel), as described elsewhere^34^. Mice in the IV cohort were intravenously administered an equal dose of PEG-840kDa-VT750. Animals were sacrificed by CO_2_ asphyxiation 60 min post-inhalation/injection and organs were removed for imaging (LICOR Odyssey). Organ fluorescence was quantified in Fiji^63^ by manually outlining organs, using the “Measure” feature, and taking the mean intensity.

Blood for pharmacokinetics measurements was collected using retro-orbital bleeds with 15 μL glass capillary collection tubes. Blood was diluted in 40 μL PBS with 5 mM EDTA to prevent clotting, centrifuged for 5 min at 5,000 × g, and fluorescent reporter concentration was quantified in 384-well plates relative to standards (LICOR Odyssey).

For immunohistochemical visualization of nanoparticles following IT administration, EZ-Link NHS-Biotin (Thermo Scientific) was coupled to PEG-8_40kDa_-amine at 2:1 molar ratio and reacted overnight, followed by spin filtration. Pulmonary delivery (50 uL volume, 10 uM concentration) was performed by intratracheal intubation. Fixation was performed 20-30 minutes later by inflating lungs with 10% formalin. Lungs were excised, fixed in 10% formalin at 4°C overnight, and embedded in paraffin blocks. 5 μm tissue slices were stained for biotin using the streptavidin-HRP ABC kit (Vector Labs) with DAB. Slides were scanned using the 20x objective of the Pannoramic 250 Flash III whole slide scanner (3DHistech).

### Mouse model and in vivo characterization

Male B6/SV129 *Kras^LSL-G12D/+^; Trp53^fl/fl^* (KP) mice between 18 and 30 weeks old were used for lung adenocarcinoma experiments. Tumors were initiated, as described previously^34^, by the intratracheal administration of 50 μL of adenovirus-SPC-Cre (2.5 × 10^8^ PFU in Opti-MEM with 10 mM CaCl_2_) under isoflurane anesthesia. Control cohorts consisted of age and sex-matched mice that did not undergo intratracheal administration of adenovirus. Tumor growth was monitored by microCT imaging (General Electric) and was scored by a blinded radiation oncologist. Each cage consisted of a combination of KP and control animals.

ABN constructs (GluFib-Substrate-PEG-8_40kDa_) for urinary experiments were synthesized by CPC Scientific (Sunnyvale, CA). ABNs were dosed (50 μL total volume, 20 μM concentration per ABN) by intratracheal intubation, as described above. All ABN experiments were performed in the morning at the Koch Institute animal facility. Bladders were voided 60 minutes after ABN administration and all urine produced 60-120 min after ABN administration was collected using custom tubes in which the animals rest upon 96-well plates that capture urine. Urine was pooled and frozen at −80°C until analysis by LC-MS/MS.

### LC-MS/MS reporter quantification

Liquid chromatography/tandem mass spectrometry was performed by Syneos Health (Princeton, NJ) using a Sciex 6500 triple quadrupole instrument. Briefly, urine samples were treated with UV irradiation to photocleave the 3-Amino-3-(2-nitro-phenyl)propionic Acid (ANP) linker and liberate the Glu-Fib reporter from residual peptide fragments. Samples were extracted by solid-phase extraction and analyzed by multiple reaction monitoring by LC-MS/MS to quantify concentration of each Glu-Fib mass variant. Analyte quantities were normalized to a spiked-in internal standard and concentrations were calculated from a standard curve using peak area ratio (PAR) to the internal standard. Mean normalization was performed on PAR values to account for mouse-to-mouse differences in ABN inhalation efficiency and urine concentration.

### Statistical analysis

For all urine experiments, PAR values were mean normalized across all reporters in a given urine sample prior to further statistical analysis. Significantly different reporters were identified by unpaired two tailed t-test followed by correction for multiple hypothesis using the Holm-Sidak method. Principal component analysis (PCA) was performed on mean normalized PAR values in the R statistical environment^61^ using the prcomp function. Binary classification was performed using the Caret package^64^ in the R statistical environment. Generalized linear model was used for RNA-seq data and random forest^50^ was used for ABN classification of urine samples. Prespecified training and validation cohorts were randomly assigned (75% training/25% validation for RNA-seq data, 50% training/50% validation for urine data). Classifiers used cross-validation on the training cohort and were trained with optimization for AUC.

## Supporting information

## Acknowledgments

We thank H. Fleming for critical editing of the manuscript; J.S. Dudani for assistance with experimental design; the KI Bioinformatics & Computing Core for assistance with binary classification and GSEA; and the KI Animal Imaging & Preclinical Testing Core for assistance with microCT imaging.

## Funding

This study was supported in part by a Koch Institute Support Grant P30-CA14051 from the National Cancer Institute (Swanson Biotechnology Center), a Core Center Grant P30-ES002109 from the National Institute of Environmental Health Sciences, the Ludwig Fund for Cancer Research, the Koch Institute Marble Center for Cancer Nanomedicine, and Johnson and Johnson. We acknowledge support from the Ludwig Center fellowship (to J.K. and A.D.W.), the National Science Foundation Graduate Research Fellowships Program (to A.D.W.), National Cancer Institute K99 Grant R00-CA187317 (to T.T.), and the Damon Runyon Postdoctoral Fellowship (to P.M.K.W.). S.N.B. and T.J. are HHMI Investigators.

## Author Contributions

J.K., A.D.W., and S.N.B. initiated and designed the study. J.K. and A.D.W. performed experiments. T.T. and P.M.K.W. generated animal models. J.C.V. performed microCT analysis. S.N.B. and T.J. supervised the research. J.K., A.D.W., and S.N.B. wrote the first draft of the manuscript. All authors contributed to writing and editing subsequent drafts of the manuscript and approved the final manuscript.

## Competing Financial Interests

J.K., A.D.W., and S.N.B. are listed as inventors on a patent application related to this work. S.N.B. is a shareholder of and consultant to Glympse Bio.

## Data and Materials Availability

The results published here are in part based upon data generated by TCGA Research Network (Fig. 2C, D, F, fig. S2; cancergenome.nih.gov), the Lung Genomics Research Consortium (fig. S2, lung-genomics.org), Chuang and colleagues (Fig. 2A, D; GSE84447)^35^, and Sweet-Cordero and colleagues (Fig. 2B, GSE49200)^36^.

