## Supplementary material for "Noninvasive lung cancer detection via pulmonary protease profiling"

Fig. S1. KP model genetically and histologically recapitulates human lung adenocarcinoma.

Fig. S2. Human LUAD-associated proteases are not overexpressed in benign lung diseases.

Fig. S3. Peptide substrates are cleaved by one or a combination of metallo, serine, and aspartic proteases.

Fig. S4. Free reporters enter the bloodstream after pulmonary delivery and are detectable in the urine by mass spectrometry.

Fig. S5. Multiple reporters are differentially enriched in the urine of healthy mice and KP mice at 7.5 and 10.5 weeks.

Table S1. Reporter and substrate sequences for *in vitro* recombinant protease screen

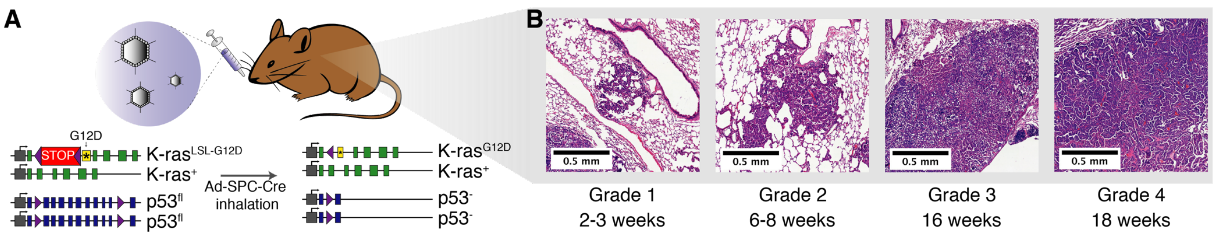


**Supplementary Fig. 1. KP model genetically and histologically recapitulates human lung adenocarcinoma.** (**A**) Disease is induced in the KP model by intratracheal instillation of adenovirus expressing Cre recombinase under the control of the SPC (surfactant protein C) promoter, which results in activation of mutant $K\text{-}ras^{G12D}$ and excision of both copies of *Trp53* in type II alveolar cells[34]. (**B**) Histologically, disease progresses from low grade dysplasia to invasive adenocarcinoma over 18-20 weeks (shown are representative lesions of each grade in a single, advanced-stage KP mouse).

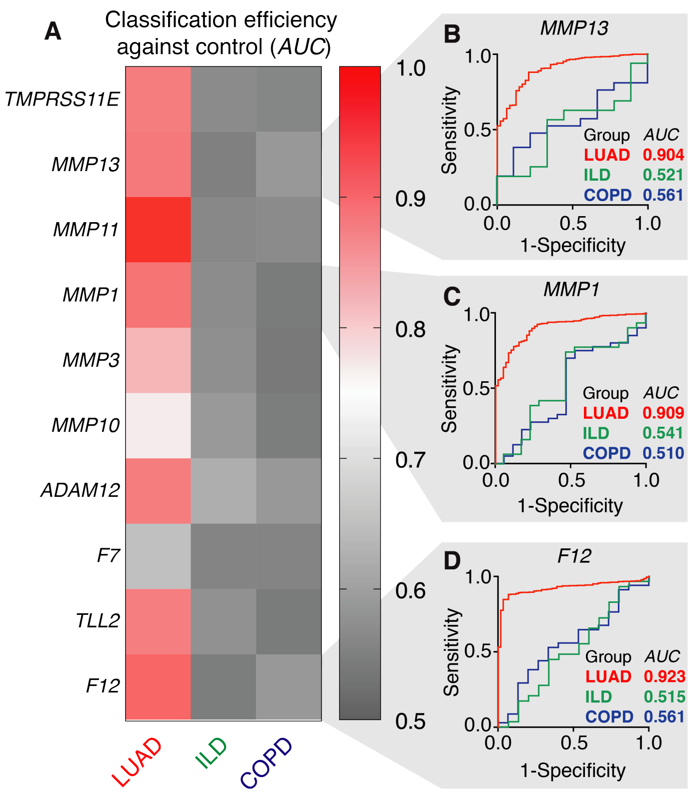


**Supplementary Fig. 2. Human LUAD-associated proteases are not overexpressed in benign lung diseases.** (**A**) RNA-Seq data curated by the Lung Genomics Research Consortium (LGRC) was analyzed to assess the classification efficiency of human lung cancer-associated proteases in interstitial lung disease (ILD, $n$ = 31) and chronic obstructive pulmonary disease (COPD, $n$ = 41) against normal lung ($n$ = 17). Of the top 20 overexpressed proteases in human LUAD, 10 were included in the LGRC dataset with FPKM values greater than zero for at least half of the samples. ROC analysis was performed for LUAD (from TCGA) and ILD and COPD (from LGRC) against their respective controls, using FPKM values for each protease. (**B-D**) ROC curves for individual proteases in the panel are shown.

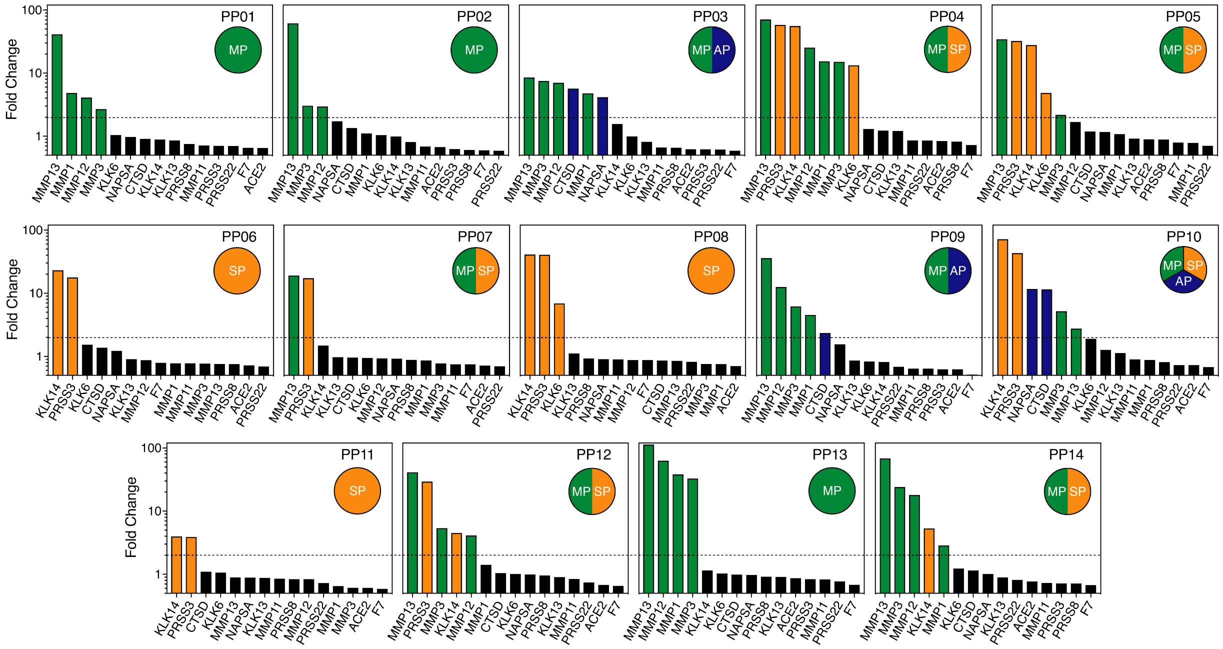


**Supplementary Fig. 3. Peptide substrates are cleaved by one or a combination of metallo, serine, and aspartic proteases.** Quantification of *in vitro* proteolytic cleavage of fluorogenic peptide substrates. Y axis represents fluorescence fold change after 45 minutes of incubation with recombinant protease and dotted line is at fold change = 2. Bars are colored according to the catalytic class of the protease (green, metalloprotease-specific; orange, serine protease-specific; blue, aspartic protease-specific).

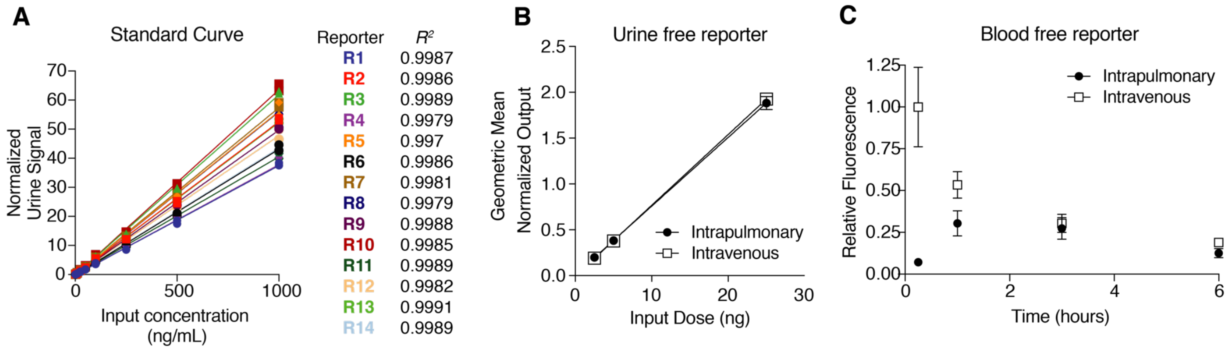


**Supplementary Fig. 4. Free reporters enter the bloodstream after pulmonary delivery and are detectable in the urine by mass spectrometry.** (**A**) Glu-fib reporters were spiked into urine at concentrations ranging from 1 to 1000 ng/mL and LC-MS/MS was performed. Goodness of fit was assessed by linear regression and is given as Pearson's $R^2$. (**B**) Healthy mice ($n$ = 4 each group) were administered MS-encoded free reporters (IT and IV) at doses ranging from 2.5 ng to 25 ng and urinary concentrations at 1 hour were assessed by LC-MS/MS ($\text{slope}_{IT} = 0.075\ \text{ng}^{-1}$, $\text{slope}_{IV} = 0.077\ \text{ng}^{-1}$). Error bars represent SD. (**C**) Cy7-labeled free reporters were administered IT and IV and concentration in the blood was assessed over the following 6 hours. Error bars represent SD.

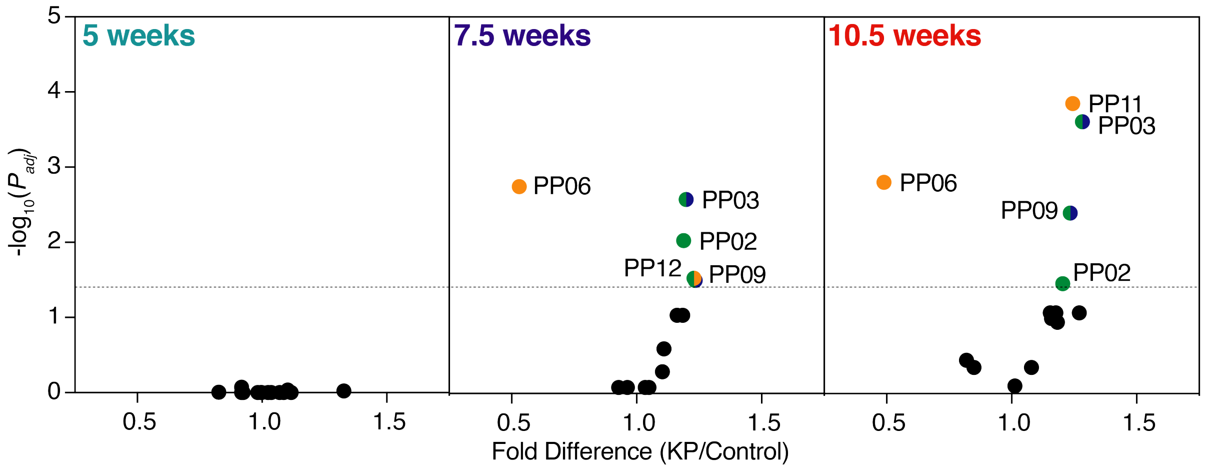


**Supplementary Fig. 5. Multiple reporters are differentially enriched in the urine of healthy mice and KP mice at 7.5 and 10.5 weeks.** Mean normalized urinary reporter concentrations in KP mice and healthy mice were compared at 5 weeks, 7.5 weeks, and 10.5 weeks after tumor initiation and $-\log_{10}(P_{adj})$ was plotted against fold difference between KP and control. Significance was calculated by two-tailed *t*-test followed by adjustment for multiple hypotheses using the Holm-Sidak method. Dotted line is at $P_{adj} = 0.05$. Significant reporters are color-coded according to the classes of protease that cleave their corresponding peptide substrates *in vitro* (fig. S3) (green, metalloprotease-specific; orange, serine protease-specific; blue, aspartic protease-specific).

| Name | Fluorophore | Substrate | Quencher |
|---|---|---|---|
| PPQ1 | 5FAM | GGPQGIWGQC | CPQ2 |
| PPQ2 | 5FAM | GGPVGLIGC | CPQ2 |
| PPQ3 | 5FAM | GGPVPLSLVMC | CPQ2 |
| PPQ4 | 5FAM | GGPLGLRSWC | CPQ2 |
| PPQ5 | 5FAM | GGPLGVRGKC | CPQ2 |
| PPQ6 | 5FAM | GGfPRSGGGC | CPQ2 |
| PPQ7 | 5FAM | GGLGPKGQTGC | CPQ2 |
| PPQ8 | 5FAM | GGGSGRSANAKGC | CPQ2 |
| PPQ9 | 5FAM | GGKPISLISSGC | CPQ2 |
| PPQ10 | 5FAM | GGILSRIVGGGC | CPQ2 |
| PPQ11 | 5FAM | GGSGSKIIGGGC | CPQ2 |
| PPQ12 | 5FAM | GGPLGMRGGC | CPQ2 |
| PPQ13 | 5FAM | GGP-(Cha)-G-Cys(Me)-HAGC | CPQ2 |
| PPQ14 | 5FAM | GGAPFEMSAGC | CPQ2 |

**Supplementary Table 1. Reporter and substrate sequences for *in vitro* recombinant protease screen.** 5FAM, 5-Carboxyfluorescein; CPQ2, quencher; Cha, 3-Cyclohexylalanine; Cys(Me), (methylsulfanyl)propanoic acid; lowercase letters, D-amino acids
